# Connecting small RNAs and Aging

**DOI:** 10.1101/021584

**Authors:** Joshua Elkington

## Abstract

**Website Summary:** Small RNAs are important gene regulators of stress response, aging, and many other things. More analysis needs to be done in order to gain a better understanding of these molecules to find connections between small RNAs and important things like aging.

**Summary:** Small RNAs are a diverse population of gene regulators, but their role in the cell is not fully characterized. Bioinformatics was used to prove their connection with aging and expand current knowledge for these molecules.

**Abstract:** Small RNAs have a wide range of functions and recent studies have found connections between these molecules and aging pathways. However, the process to systematically characterize this relationship is slow. Prediction tools can be used to expedite this process by finding new genes and pathways that cross talk with each other. Using phylogenetic and systems analysis, connections between small RNAs and aging were proven and new genes that may be related to aging were identified. This type of analysis can be applied to many different pathways in order to fully characterize the role of small RNAs.

## Introduction

The phenomenon of RNAi interference (RNAi) was first found in the nematode, *C. elegans*. In RNAi, dsRNA is processed to silence homologous genes. The discovery of small RNAs in worms has led to the discovery of thousands of endogenous small RNAs that fall into three categories: microRNAs, endogenous small interfering RNA (endo-siRNAs), and Piwi-interacting RNAs (piRNAs)^1,2,3^.

These small RNAs target coding genes, transposons, and pseudogenes to regulate a wide range of pathways through Argonuate proteins. For example, piRNAs interact with PRG-1/2 and endo-siRNAs bind ERGO-1.

Of the three classes of small RNAs, endo-siRNAs remain the least well understood. Endo-siRNAs have a 5’ guanosine that either are 26 nucleotides (nt) long or 22 nt^4,5,6^. The 26G and 22G endo-siRNAs have overlapping biogenesis components but engage distinct pathways. The 22G RNAs bind worm-specific Argonautes (WAGO) and the Argonaute, CSR-1^7,8^. Approximately half of the 27 Argonaute proteins encoded in *C. elegans* belong to a WAGO clade.

microRNAs have been found to regulate diverse pathways related to development, stress response, and longevity. However, endo-siRNAs and piRNAs are thought to maintain the germline of worms by genome surveillance. Recently, genes that specifically regulate endo-siRNAs have been discovered. From these genes, *eri-6/7, alg-3/4* and *ergo-1* are especially interesting due to their possible connections to aging. ERI-6/7 is a helicase protein required for ERGO-1 dependent 26G and 22G RNA accumulation^9^. The ERGO-1 Argonaute protein binds and stabilizes 26G RNAs in the germline^10,11^. ALG-3 and ALG-4 are homologous Argonautes that bind and stabilize 26G RNAs in spermatogenic germline. Both *eri-6/7* and *ergo-1* exhibit an enhanced RNAi phenotype associated with loss of 26G RNAs, and ERI-6/7 is thought to function in Argonaute loading of RNA^12^ (Figure 1). These proteins related to endo-siRNAs help maintain the integrity of the genome, and as a result, this pathway may be involved in regulate lifespan because of the connection between longevity and viability.

**Figure 1.**
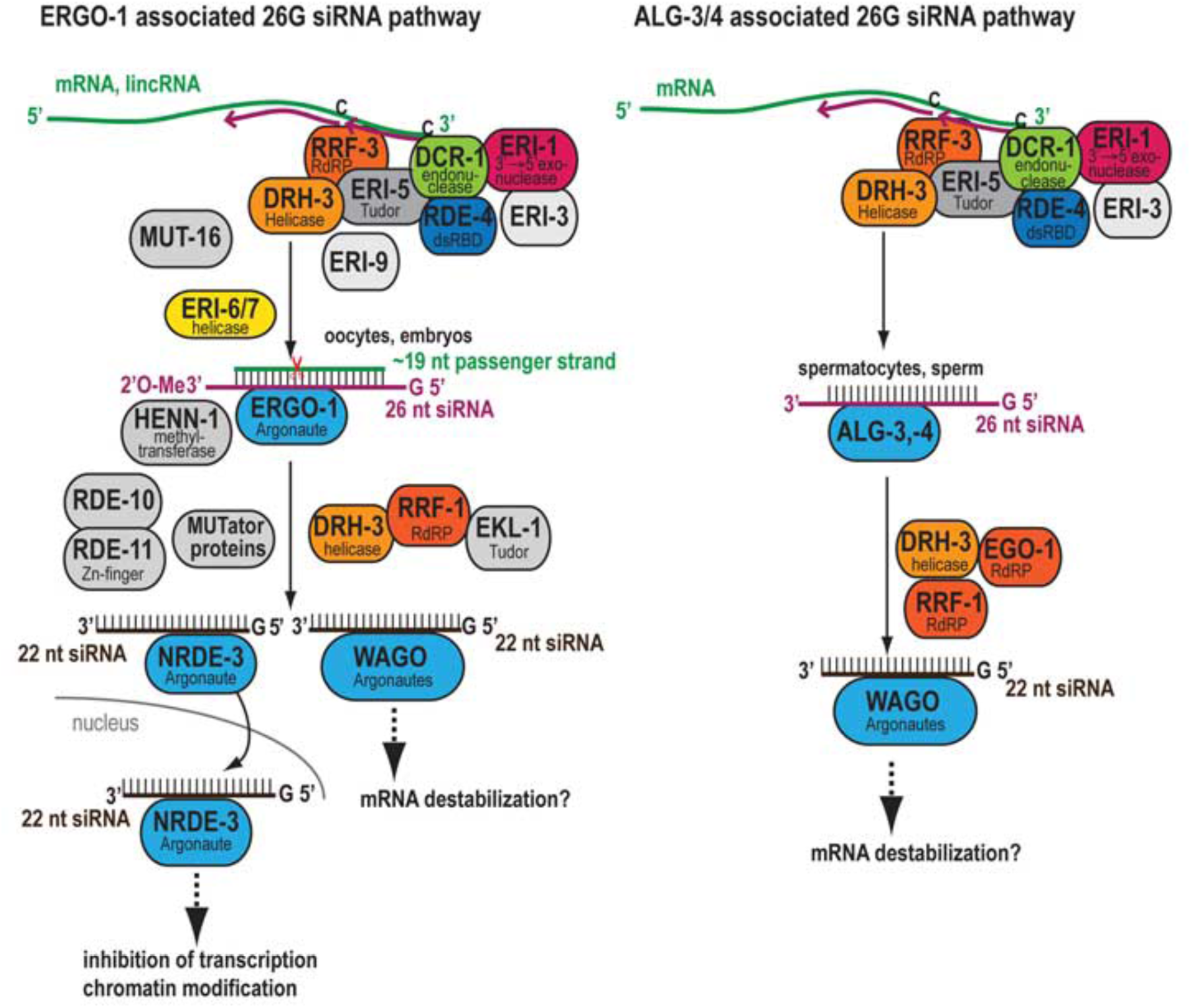
26G endo-siRNA pathways. A complex wit DCR_1 generates 26G endo- siRNAs from mRNA and lncRNA templates. The processed siRNAs interact with the Argonautes, ERGO-1 and ALG-3/4. The generation of 22G siRNAs use the mRNA template bound by 26G siRNAs and require unique genes for amplification. The 22G siRNAs interact with WAGO Argonautes.

The widely conserved *C. elegans* insulin/IGF-1 signaling (IIS) pathway regulates longevity, metabolism, growth, development, and behavior. This pathway is activated by insulin-like ligands that bind the insulin/IGF-1 transmembrane receptor (IGFR) ortholog, DAF-2. This receptor controls a kinase cascade that regulates a FoxO transcription factor, DAF-16 that regulates genes related to stress response^13,14^(Figure 2).

**Figure 2.**
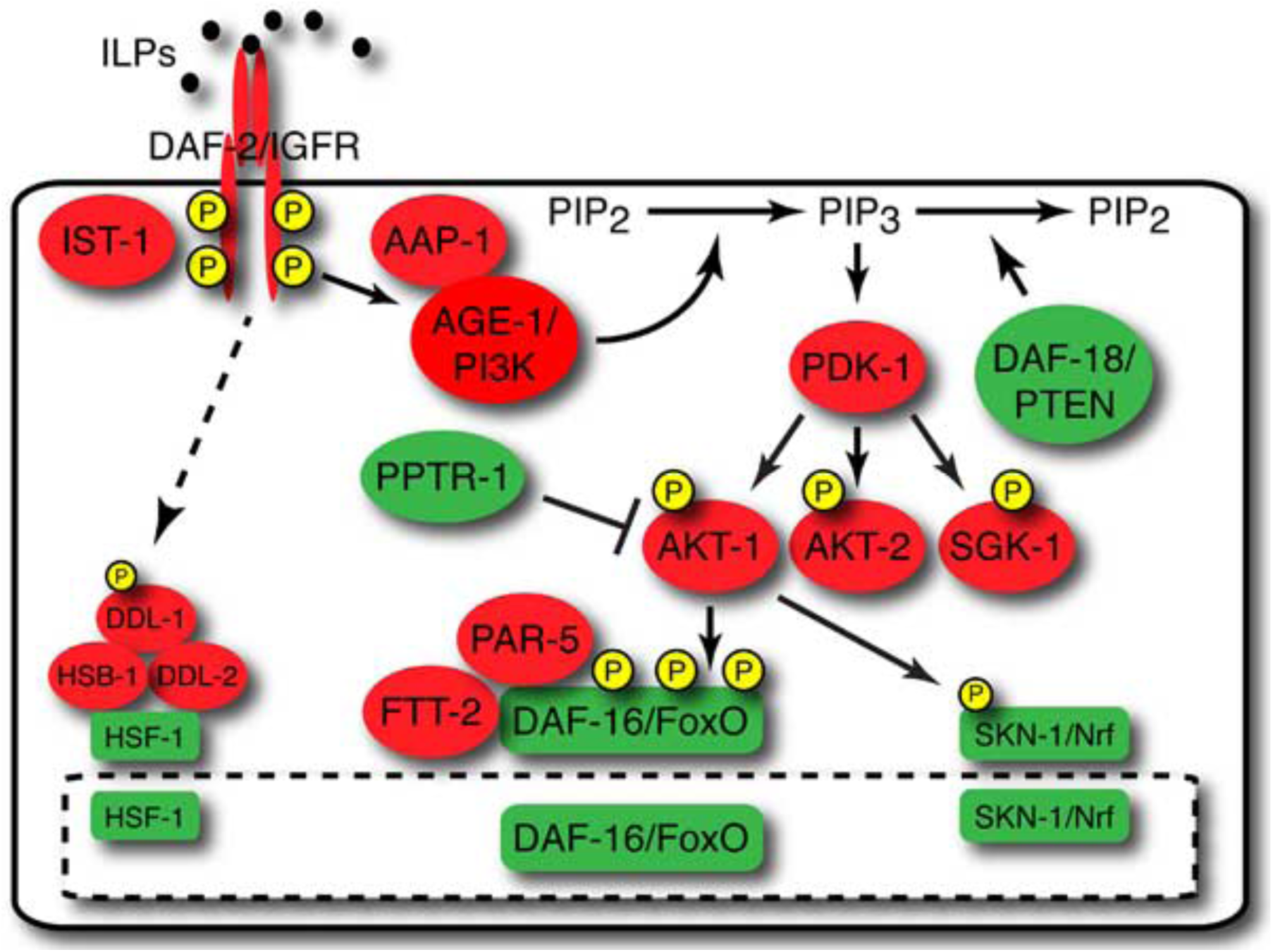
*C. elegans* Insulin Pathway. Activation of the DAF-2 receptor by an insulin-like ligand promotes phosphorylation of DAF-2 and SKN-1 to prevent entry into the nucleus.

Recent research has proven a connection between the IIS pathway and piRNAs that implies that aging and transgenerational maintenance of germ cell are connected. *daf-2* mutants inhibited *prg*-dependent sterility^15^. Starved *prg-1* mutants had an extended lifespan that was dependent on *daf-16,* and silencing of repetitive loci was restored in *prg-1;daf-*2 mutants^15^. These recent findings prove that piRNAs and IIS are related, and provide evidence that small RNA pathways act through pathways related to aging. Therefore, there may be a connection between endo- siRNAs and components of the IIS pathway. Using phylogenetic analysis and gene interactome data, connections between the insulin pathway and endo-siRNAs pathway.

To confirm that this analysis is valid, prg-1/2 and genes related to aging are found to co-evolve with one another and share overlapping interacting partners.Next, this analysis was applied to ergo-1 and eri-7 in order to prove the connection of endo-siRNAs with the IIS pathway and aging.

## Overview

### Phylogenetic Analysis

Phylogenetic analysis was used to determine genes that co-evolve with one another^16^. BLAST scores of protein genes in *C. elegans* were normalized to the length of the query sequence and relative phylogenetic distance from *C. elegans* for each of the 86 organism genomes. For the 10,054 *C. elegans* proteins that have paralogs in other organisms, single proteins were queried to generate a ranking of other *C. elegans* proteins that have the most similar pattern of conservation values. Proteins that are in the same families have similar patterns of conservation across evolution and cluster together by this analysis. More importantly, proteins that have no apparent similarity cluster together. For the genes discussed, the top 50 genes that co-evolved with them were used for further analysis.

### Genetic System Analysis

GeneMANIA was used in order to prove that genes from the phylogenetic analysis are related and find new members of this network. This tool finds other genes that are related from a set of input genes. GeneMANIA used a large set of functional associated data such as protein and genetic interactions, pathway knowledge, co-expression, co-localization, and protein domain similarity. This tool can be used to enrich for new components of a known pathway or molecular complex. GeneMANIA expands that genetic network determined by the phylogenetic analysis. This method was used to analyze genes that co-evolved with prg-1/2 and eri-7 and ergo-1.

## Results

### piRNAs and aging

Using phylogenetic and genetic system analysis, PRG-1/2 is related to the aging gens DAF-16 and PHA-4. PHA-4, UNC-130, and K04C1.3 co-evolved with PRG- 1/2 (Figure 3, 4). By analyzing the interaction system, DAF-16 was fond to interact wit UNC-130, and K04C1.3, and PHA-4. Furthermore, ALG-2 was found to interact with PHA-4 and co-evolve with PRG-1/2 (Figure 5).

**Figure 3.**
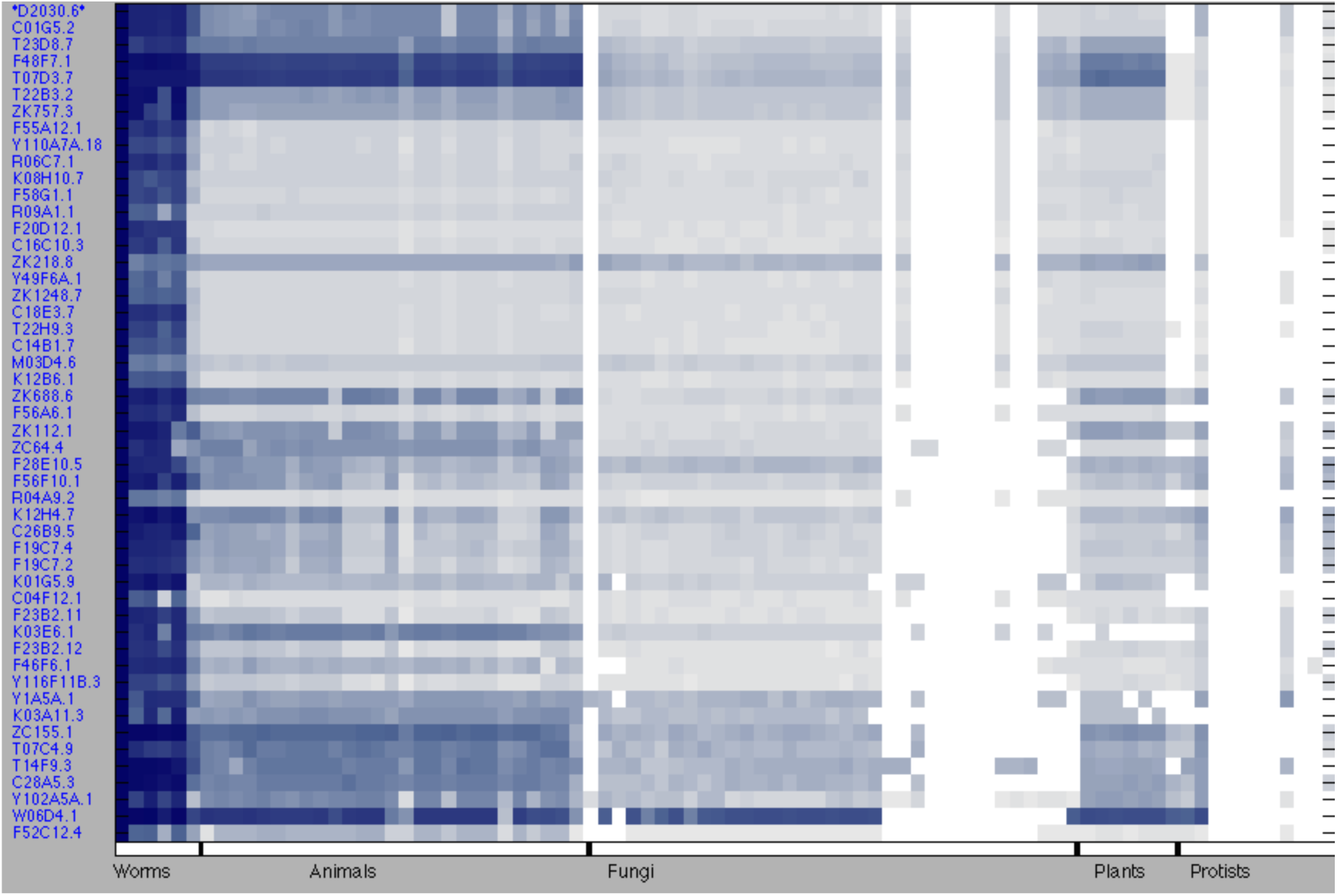
PRG-1 Phylogenetic profile of PRG-1.

**Figure 4.**
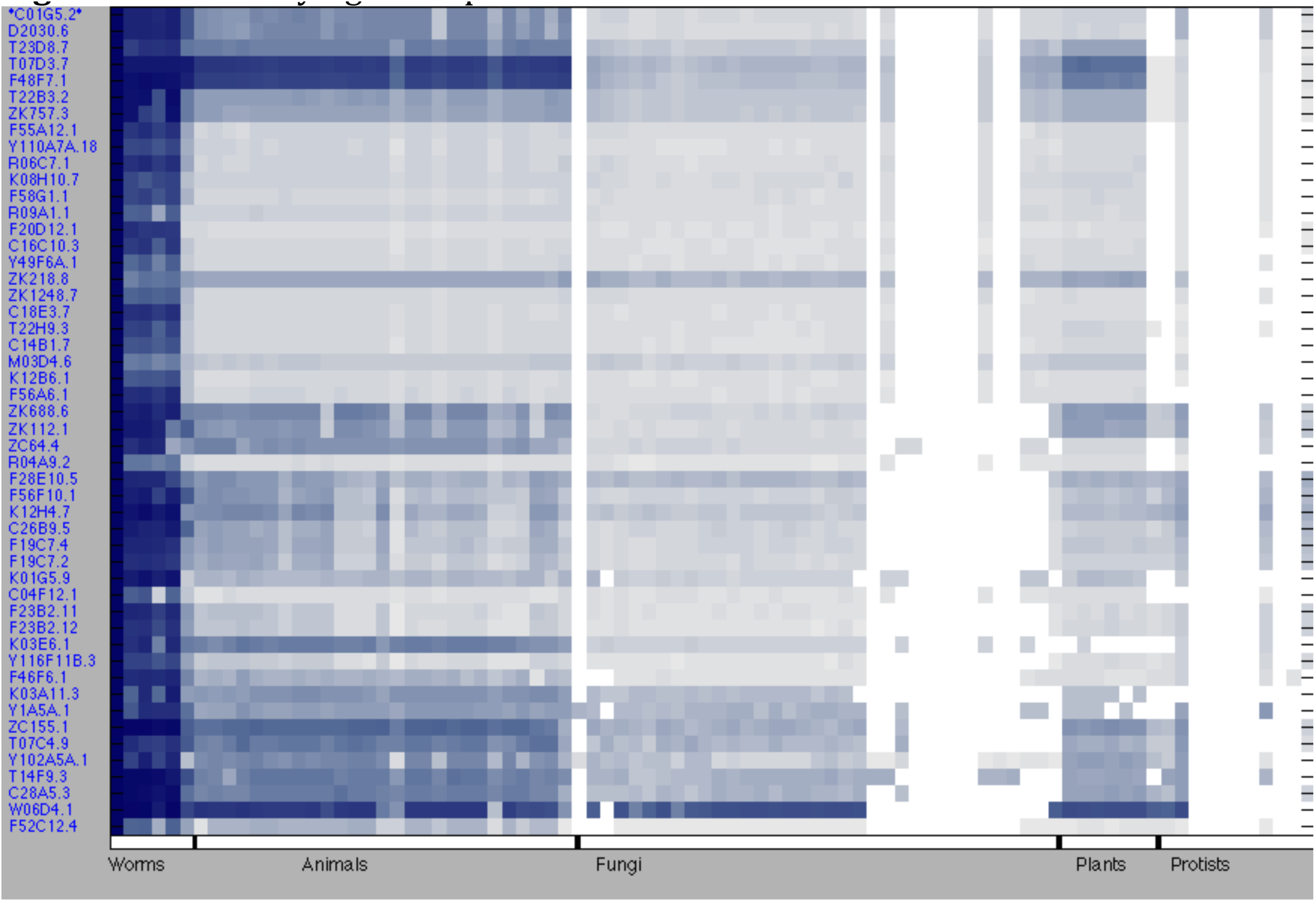
PRG-2. Phylogenetic profile of PRG-2

**Figure 5.**
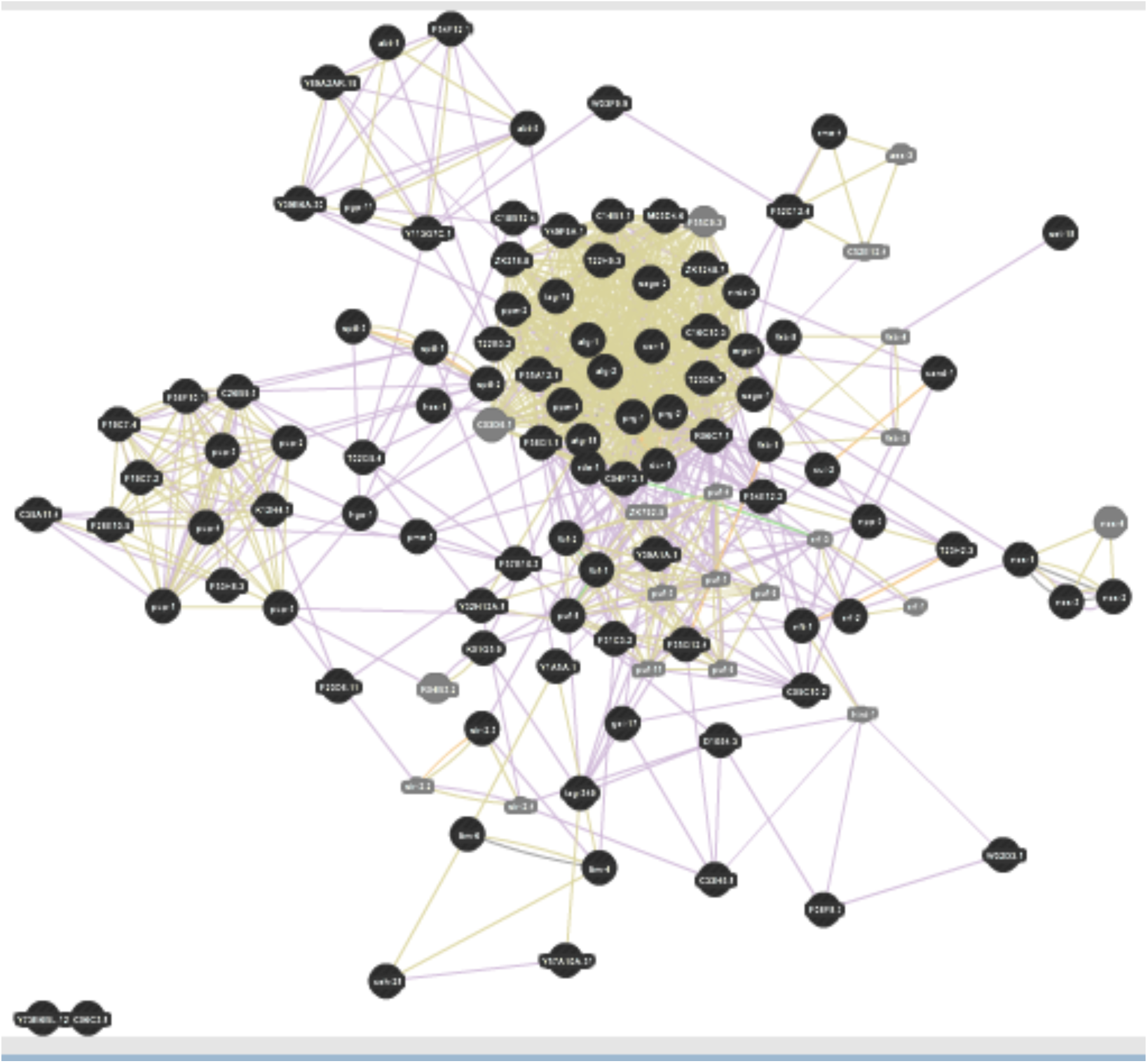
PRG-1/2 System. Genetic interaction system of genes that co-evolve with PRG-1 and PRG-2.

### Endo-siRNAs and aging

Many genes related to small RNA pathways were found to co-evolve with ERGO-1 and ERI-7 (Figure 6, 7). After doing systems analysis on the set of genes, many genes related to aging were enriched. DAF-16, SKN-1, AGE-1, and PHA-4 were found to interact with many genes that co-evolved with ERGO-1 and ERI-7 (Figure 8).

**Figure 6.**
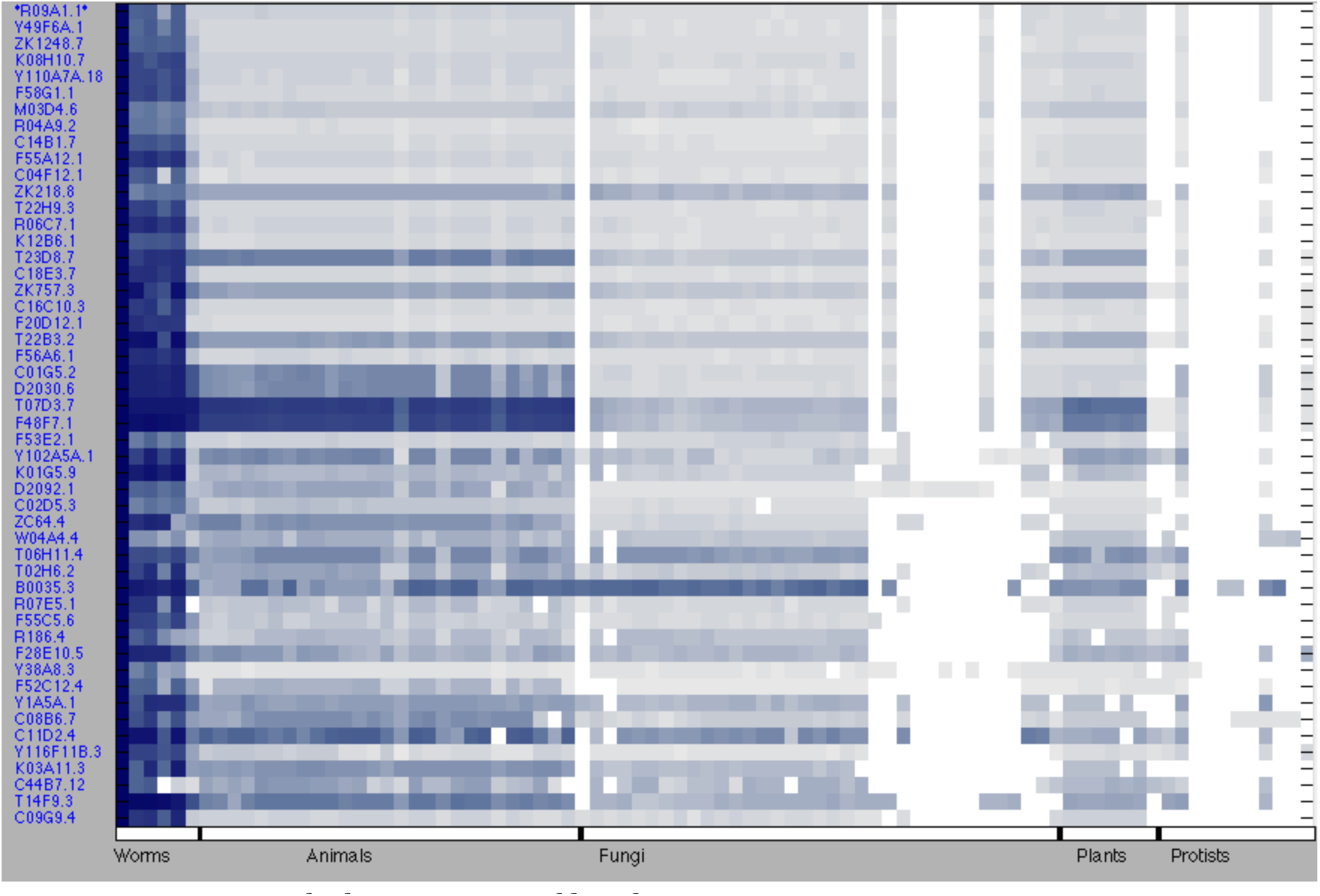
ERGO-1. Phylogenetic profile of ERGO-1

**Fig 7.**
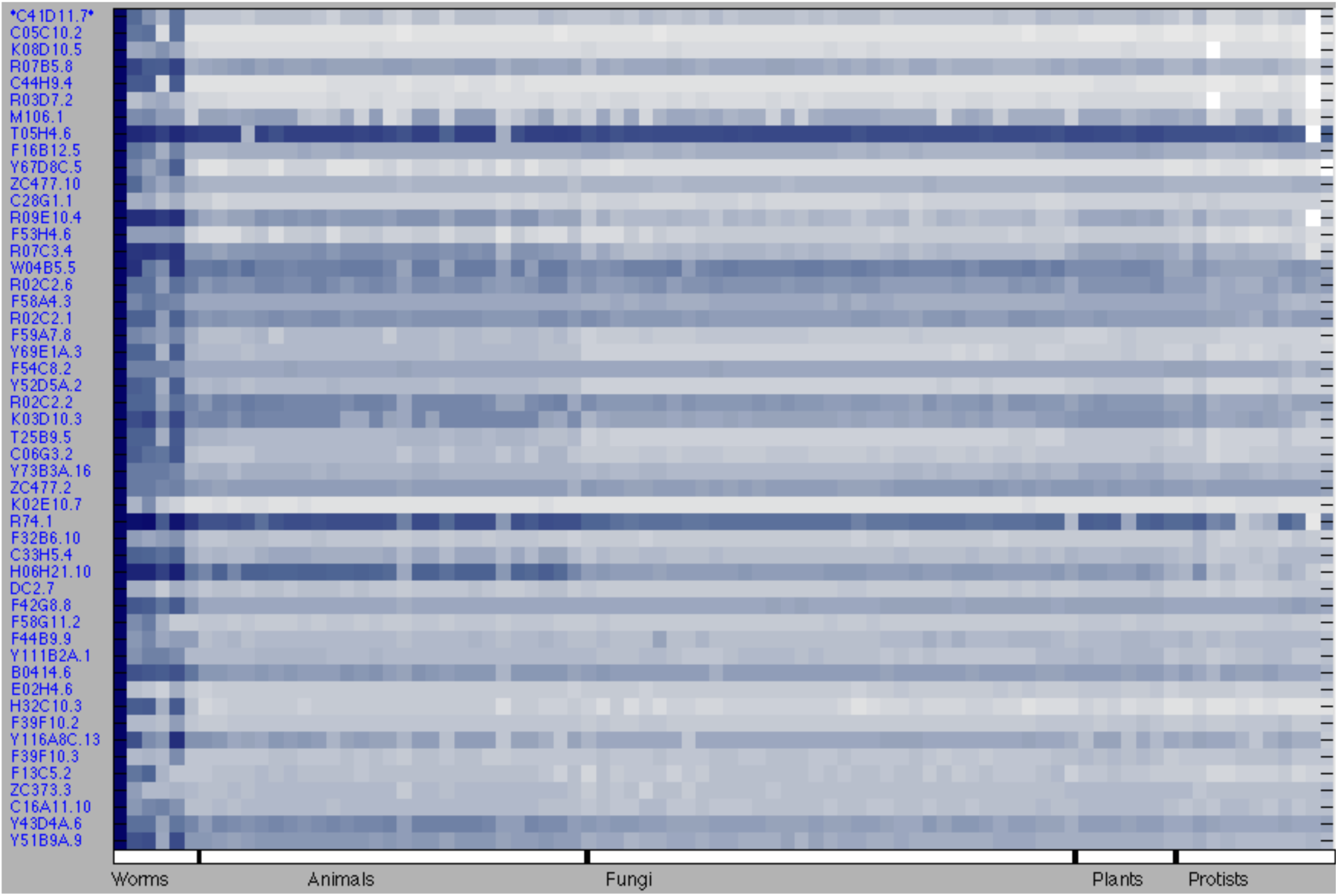
Figure 7 ERI-7. Phylogenetic profile of ERI-7

**Figure 8.**
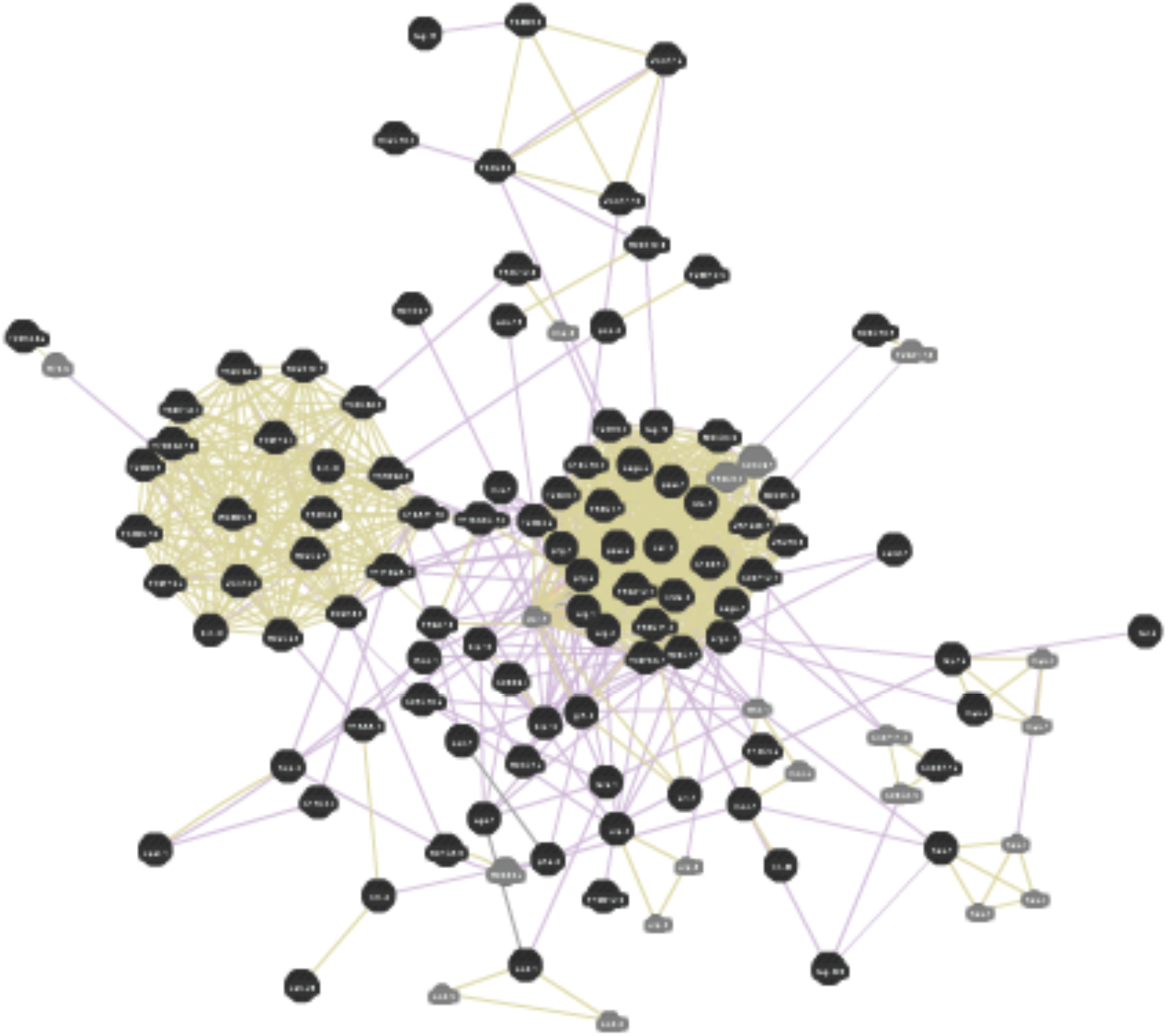
ERGO-1 and ERI-7 System. Genetic interaction system of genes that coevolve with ERGO-1 and ERI-7.

## Discussion

### Evolutionary role of small RNAs

Small RNAs are a diverse set of molecules that regulate a wide range of pathways. Ultimately, small RNAs are involved in the maintenance of organism integrity. The enrichment of piRNAs and endo-siRNAs in the C. *elegans* germline and their role in genome surveillance would make them good candidates to be involved in regulating longevity. Recent findings have connected piRNAs with the IIS pathway, and this analysis shows that piRNAs and endo-siRNAs are related to aging- related genes. As a result, through evolution these classes of small RNAs may have adopted significant roles in regulating aging along with their functions in maintaining genome integrity.

### Relationship between aging and small RNAs

The connections between aging and small RNA pathways are not well understood. Using prediction tools like phylogenetic analysis can help guide experiments to characterize this relationship. So a next step to understand the role of small RNAs in aging is to prove that the genes predicted to connect these two pathways significantly connect two pathways. Maybe induction of stress or a longevity signal could activate a cascade that up regulate small RNAs in order to set a threshold on gene expression to adapt to the environment. Possibly small RNAs could directly regulate aging genes or they could receive environmental signals themselves in order to modulate organism lifespan through existing or new aging pathways.

### Power of phylogenetic profiling

Phylogenetic profiling is a unique tool that can find proteins with similar patters of conservation and divergence across many organisms. Proteins that have similar phylogenetic patterns tend to function in the same pathways. Protein divergence is not a random event because entire classes of proteins are lost together in particular taxa. As these organisms specialize, entire classes of proteins are gained and lost together. Analyzing the evolutionary relationship of genomes of many different organisms gives insights into new relationships between known pathways.

